# Gaussian decomposition of high-resolution melt curve derivatives for measuring genome-editing efficiency

**DOI:** 10.1101/176719

**Authors:** Michail Zaboikin, Carl Freter, Narasimhachar Srinivasakumar

## Abstract

We describe a method for measuring genome editing efficiency from *in silico* analysis of high-resolution melt curve data. The melt curve data derived from amplicons of genome-edited or unmodified target sites were processed to remove the background fluorescent signal emanating from free fluorophore and then corrected for temperature-dependent quenching of fluorescence of double-stranded DNA-bound fluorophore. Corrected data were normalized and numerically differentiated to obtain the first derivatives of the melt curves. These were then mathematically modeled as a sum or superposition of minimal number of Gaussian components. Using Gaussian parameters determined by modeling of melt curve derivatives of unedited samples, we were able to model melt curve derivatives from genetically altered target sites where the mutant population could be accommodated using an additional Gaussian component. From this, the proportion contributed by the mutant component in the target region amplicon could be accurately determined. Mutant component computations compared well with the mutant frequency determination from next generation sequencing data. The results were also consistent with our earlier studies that used difference curve areas from high-resolution melt curves for determining the efficiency of genome-editing reagents. The advantage of the described method is that it does not require calibration curves to estimate proportion of mutants in amplicons of genome-edited target sites.

## Introduction

Genome editing at predetermined loci has been greatly facilitated by new technologies based on RNA-guided endonucleases (RGENs)[1–3] or transcription-activator like effector nucleases (TALENs) [4–6]. The sequence-directed endonucleases introduce doublestranded breaks (DSBs) at the target site. The DSBs can undergo two major types of DNA repair. Non homologous end joining (NHEJ) repair results in indels at the cut site. Homology-directed repair (HDR) either restores the original in the presence of an endogenous template (sister chromatid) or inserts an exogenous DNA donor template when available across the cut site [7–9].

The ability to generate genome-editing reagents with a desired specificity does not guarantee efficient target site modification. There is therefore a need for methods that rapidly assess reagent efficacy. A common approach is to determine efficacy of genome editing reagents is to transfect human embryonic kidney (HEK293T) cell line with the reagents. This is followed by amplification of target region by PCR and generation of heteroduplexes by denaturation and renaturation in the presence of unmodified wild type or different alleles. Mismatches in these heteroduplexes can be identified by digestion with single-strand specific endonucleases (such as T7 or Surveyor nuclease) and resolution of the digestion products in polyacrylamide or agarose gels [10–12].

A second approach to determine efficacy of genome editing is to use TaqMan assays with probes designed to bind over the putative target cut site [12,13]. Reduced binding of the TaqMan probe, due to indel mutations at the target site, with reference to a control TaqMan probe that binds outside the cut site, can be used to estimate the editing efficacy.

A third method, which is gaining popularity, uses high resolution melting analysis (HRMA) after real-time PCR with nonspecific double-stranded DNA (dsDNA)-binding dyes such as Eva Green [12,14–16]. These dyes are more fluorescent when bound to dsDNA. In this method, after amplifying the target region containing the repaired double-stranded break site, the dsDNA is gradually warmed until the DNA completely melts. As dsDNA regions melt into single-stranded regions, dye is expelled, decreasing the fluorescence signal. Melting characteristics depend on the length of the PCR product, the sequence, and the GC content. The temperature at which half of the DNA is single-stranded is called the Tm. The Tm peak can be readily identified by first derivative transformations of melt curve data. Target cut sites repaired by NHEJ generally exhibit lower Tms as the amplicons are usually of smaller size than the wildtype target PCR product. We previously used HRMA to estimate RGEN editing efficiency [12]. In that study, the region encompassing the target site was amplified in a real-time PCR buffer and subjected to HRMA. Normalized melt curves from genome-edited test samples were subtracted from control curves obtained from unmodified targets to obtain difference curves. The difference curve areas (DCAs) related directly to the percentage of mutants in the PCR product. We used standard curves generated with mixes of wild type and mutant PCR products to accurately estimate the percentage of mutants in different test samples. A major bottleneck to this method was the requirement for a purely mutant PCR product to generate mixes for calibration curves.

Here we describe an alternative method that does not require standard curves to measure the proportion of mutant species from high-resolution melt curve data. The high resolution melt curves were first corrected for temperature dependent quenching of free and ds-DNA bound fluorophore and then numerically differentiated to obtain first derivative melt curves. First derivative melt curves from unmodified control target sites were modeled as sum of two Gaussian components while edited samples were modeled using an additional Gaussian component for the mutant population discernible in first derivative melt curves. The weight of the “mutant” Gaussian component was shown to accurately reflect editing efficiency of sequence-directed endonucleases.

## Materials & Methods

### Cells

Human embryonic kidney (HEK293T) cells were maintained in Dulbecco’s modified Eagle’s medium containing 2 mM L-glutamine, 100 U/ml of penicillin, 100 μg/ml streptomycin and 10% heat-inactivated fetal bovine serum (FBS) (Hyclone/ThermoFisherScientific, USA) as described previously [17,18].

### Plasmids

The plasmid constructs encoding TALENs targeting the c-c motif chemokine receptor 5 (CCR5, GenBank RefSeqGene number NG_012637) intron immediately downstream of the coding exon have been described [12]. The dimeric guide RNA (dgRNA)-dCas9-FokI system consists of pSQT1313 and pSQT1601 plasmids. pSQT1313 is used for expression of dual guide RNAs (gRNAs) that target genomic DNA sequences on opposite strands and spaced approximately 16 bases apart. pSQT1601 encodes dCas9-FokI fusion protein to effect DSBs and Csy4 RNase to process the dgRNA expressed by pSQT1313. The dgRNA-dCas9-FokI system was a gift from Keith Joung via Addgene.org. pSQT1313-F8S2, targets the human coagulation factor VIII (F8) intron site 2 (F8-S2) and has been previously described. The targeting/donor plasmid (pDonor-F8) or its backbone construct (pBackbone) have also been described previously and encode a drug-resistance marker that allows selecting transfected cells using puromycin.

### CaPO_4_-mediated transfection

Plasmids were introduced into sub confluent cultures of HEK293T cells in 6-well plates by CaPO_4_-mediated transient transfection protocol as described previously [18]. Following transfection, genomic DNA (gDNA) was isolated from unselected or puromycin-selected populations using Qiagen DNeasy Blood and Tissue kit (Qiagen, Maryland, USA) as per the recommended protocol.

### Amplification of target loci for obtaining high-resolution melt curves

This has been detailed in our earlier study [12]. Briefly, gDNA from genome-edited samples was amplified using primer pairs SK144 and SK145 for the CCR5 locus, and SK228 and SK229 for the F8-S2 locus, in Precision Melt buffer (Bio-Rad, USA). SK144 and SK145 generate a PCR product of size 107 bp. For some experiments we used a different forward primer, SK214, that was located further upstream and produced a PCR product of size 140 bp with reverse primer SK145. The sequences and genome locations of these primers have been described earlier [12]. The gDNA from unmodified or mock-transfected cells were also amplified in parallel using the same primer pairs. Post amplification melting of the PCR product was done between 65°C to 95°C in 0.2°C increments.

### Processing melt curve data

Relative fluorescence units (RFUs) of melt curve data were processed to correct for background fluorescence of “unbound” fluorophore and for the temperature-dependent quenching of dsDNA-bound fluorophore as described below.

For background fluorescence correction of unprocessed RFU, we used the post-melt region of individual melt curves identified from plots of the raw RFU vs. temperature. We plotted this region separately to obtain the parameters of a linear least squares fitting. From this equation, we were able to extrapolate the background RFU at each of the measured temperature points (Equation 1). Subtracting this value from the raw RFU gave us the background subtracted RFU (BcRFU) (Equation 2).

The equations for background fluorescence correction of raw RFU:

Extrapolation of post-melt region using a first-order polynomial,

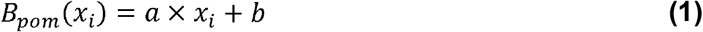

where, *x*=temperature (°C) and *T_low_ ≤ x_i_ ≤ T_high_*
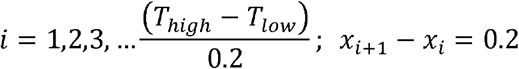 (temperature increment unit)
*T_low_* and *T_high_* refer to the lower (e.g., 71°C) and higher (e.g., 95°C) limits of the temperature range selected for melt curve analysis The slope “*a*”, and the y-intercept “*b*” parameters are obtained from first-order polynomial least-squares fitting of the post-melt region of the melt curve.

Background subtracted RFU,

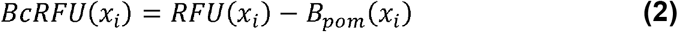

The pre-melt region of a melting curve identified from plots of melt curves of unmodified or mock-transfected cells was used to determine the efficiency of detecting dsDNA-bound fluorophore at different temperatures. This region of *BcRFU(x)* of mock-transfected cells was plotted separately and subjected to least squares curve fitting (Equation 3). The curve-fitting equation was then used to extrapolate the values across the entire range of temperatures encompassing the melting curve. The resulting values, representing predicted RFU of unmelted DNA at the different temperatures, were then normalized to the starting temperature (*T*_low_ or 71°C) to obtain the efficiency of detection of dsDNA-bound fluorophore at each measured temperature point (Equation 4). The detection efficiency of dsDNA-bound fluorophore derived from multiple mocks were averaged. The *BcRFU(x)* of mock or test samples were then divided by the average efficiency to obtain unquenched or fluorescence-corrected RFU (*FcRFU(x)*) at each temperature point (Equation 5). The *FcRFU(x)* at *T*_low_ (71°C) was then used to normalize the melt curve to yield normalized *FcRFU(x)* or *nFcRFU(x)* (Equation 6). First derivatives of nFcRFU, obtained by numerical differentiation (Equation 7), were used for subsequent curve fitting analysis.

The mathematical formulations for correction of *BcRFU(x)* for temperature-dependent quenching of fluorescence of dsDNA-bound fluorophore are shown below.

Extrapolation of pre-melt region,

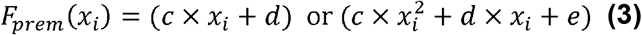

where, the parameters *c, d, e* were obtained from 1*^st^*-
 or 2*^st^*-order polynomial least squares fitting
 of pre-melt region of *BCRFU(x)*

Efficiency of dsDNA detection at temperature

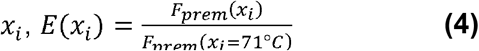

Fluorescence corrected-RFU,

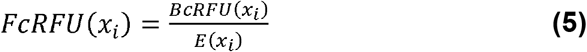

Normalized FcRFU,

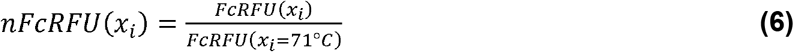

(where *nFcRFU(x_i_)* represents dsDNA content ranging from 1 in the pre-melt region to 0 in post-melt region)

The numerical differentiation of *nFcRFU(x)* was carried out as follows:

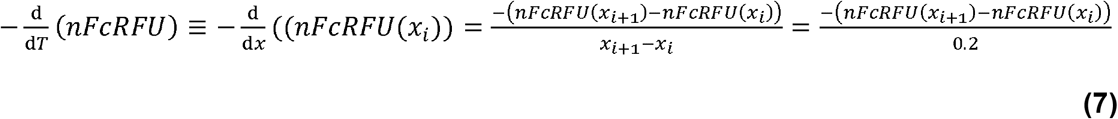

### Gaussian decomposition of first derivatives melt curves of unedited control samples

Gaussian decomposition (GD) of first derivatives of *nFcRFU(x)* was done using a commercial software, CurveExpert Professional (V. 2.6, created by Daniel Hyams, Madison, AL, USA). The normalized melt curve spans between zero and one and resembles a cumulative probability distribution function. The first derivative of the normalized melt curve resembles the density of probability distribution. A normal density distribution is mathematically represented as:

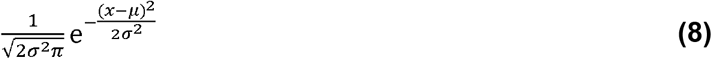

where, µ is the center of the peak, σ is the standard deviation or SD (width at half-maximal height of peak) and *x* is the temperature variable. For simplicity, we refer to this function hereafter as Gaussian function or Gaussian in place of the more cumbersome “probability density of normal distribution”.

Since, the actual Gaussian function is of the form 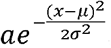, *a* corresponds to 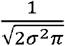 in Equation 8 where the probability density distribution has been integrated and normalized to one (the area under the curve).

For Gaussian modeling of derivative melt curves from unmodified control samples, the first derivate of nFcRFU from mock-transfected (unmodified loci) samples were modeled as either a single Gaussian function, *g2(x)*:

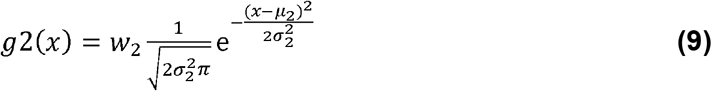

where, the free parameter *w*_2_ represents the area under the curve or weight.
or as the sum of two Gaussian components, *g2(x)* and *g3(x)*:

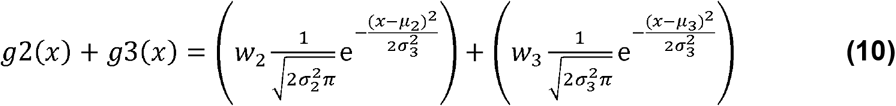

where, the Gaussian weights, *w*_2_ + *w*_3_ =1 or *w*_3_ =1–*w*_2_.

The parameters *µ_2_*, and *µ_3_*, refer to the peak center or mean, and *σ*_2_ and *σ*_3_ refer to the corresponding standard deviations (SDs) of Gaussian functions *g2(x)* and *g3(x)*, respectively. From curve fitting using the sum of two Gaussian functions (*g2(x)* and *g3(x)*), we were able to determine and ‘fix’ the parameters *w_2_*, *w_3_*, *µ_2_*, and *µ_3_* for subsequent determination of percentage of mutants in genome-edited test samples (see below).

### GD of genome-edited samples

For GD of derivative melt curves from genome-edited samples, the first derivative of *nFcRFU(x)* from test samples with genome-edited target loci were curve fitted as a sum of either two Gaussian functions, *g1(x)* and *g2(x)* or as the sum of three Gaussian functions, *g1(x)*, *g2(x)* and *g3(x),* where *g1(x)* represents the contribution of the mutant population, and *g2(x)* and *g3(x)* representing the contribution of the wildtype population in the PCR amplicon of a given target site.

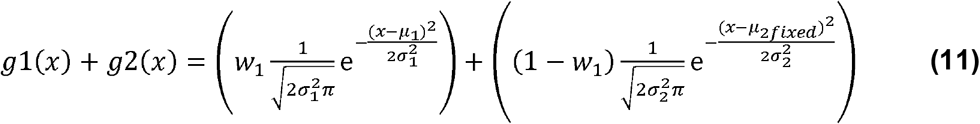

where, *w_1_ + w_2_ = 1*; the ‘fixed’ parameter µ_2fixed_ was determined from curve fitting of mock samples using the single-Gaussian function, *g2(x)*, the other parameters were set free.

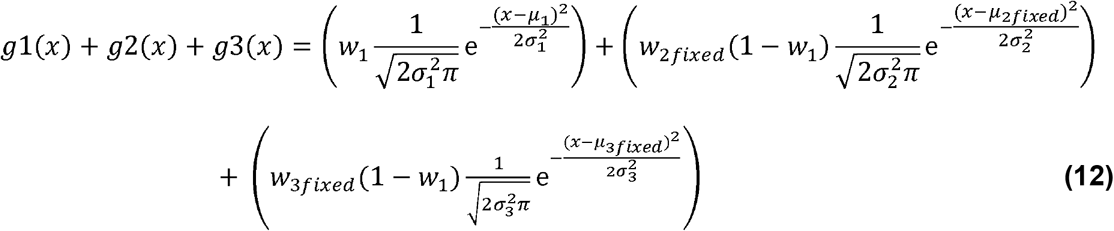

where, *w_1_ + w_2fixed_(1-w_1_) + w_3fixed_(1-w1) = 1*, and *w_2fixed_*, *w_3fixed_*, *µ_2fixed_*, and *µ_3fixed_* were determined from curve fitting of mock samples as the sum of two Gaussian functions, *g2(x)* and *g3(x)*, the other parameters were set free. The *w_1_* parameter determined from curve fitting using either *g1(x) + g2(x)* or *g1(x) + g2(x) + g3(x)* functions represents the mutant frequency in the amplicon.

### Curve fitting model comparison

CurveExpert Professional outputs the corrected Akaike Information Criteria (AICc) values for comparing curve fitting models – the model with the lower AICc value is deemed to have the better fit. The relative likelihood was calculated using *e^-0.5×(AICc_min_-AIC_ci_)^* where AICc_min_ is the model with the lower of the two values and AICci is the value of the alternate model. CurveExpert Professional also provides fitting “scores” for models, ranging from zero to 1,000 with a higher score indicating a better fit. The score is in part based on Akaike information criteria (AICc). The CurveExpert Professional scores were compared using Student’s t-test (paired, two-tailed).

## Results

### High-resolution melt curve analysis

The high-resolution melt curve data used here were generated in an earlier study [12]. Briefly, HEK293T were transfected with genome-editing reagents using a CaPO_4_ method. Two target regions were edited: F8 intron 1, and the CCR5 intron immediately downstream of the coding exon. Although we targeted three distinct sites within the F8 intron in the earlier study (referred to as sites F8-S1, -S2 or -S3), here we use data from genome-edited F8-S2 only. We used TALENs for editing the CCR5 locus and dgRNA/dCas9-FokI based RGEN system for editing the F8-S2 site. The gDNA, isolated from unselected or selected populations of transfected cells, were amplified and high-resolution melt curve data were obtained as described in Materials and Methods.

A high-resolution dsDNA melting curve consists of three regions: An initial pre-melt region where the DNA is double-stranded, followed by a transition to more rapid decrease in fluorescence attributable to DNA melting (melt region), and a second transition to a postmelt region where the DNA strands are fully separated. The pre-melt region exhibits a downward or negative slope with an increase of temperature prior to the transition to melting. This decrease in fluorescence of dsDNA-bound fluorophore prior to the beginning of separation of DNA strands can be attributed to temperature-dependent quenching of fluorescence of dsDNA-bound fluorophore. The post-melt region also exhibits a downward slope, albeit much shallower than the pre-melt slope. Since the post-melt region should contain only unbound or free fluorophore, the decrease seen in this region can be attributed to quenching effect of temperature on free or unbound fluorophore. Even after correcting melt curve data for these two quenching phenomena, the resultant melting curves of different samples frequently exhibit different pre-melt (starting) RFUs necessitating a normalization step. The raw fluorescence, reported as relative fluorescence units or RFU, therefore require processing and normalizing to enable comparison of different melting curves and for decomposition into their Gaussian components.

### Correction of RFU for temperature-dependent quenching of free fluorophore

To mathematically approximate free fluorophore behavior in the post-melt region, and to determine the effect of temperature on fluorescence of free fluorophore over the entire temperature range of melting, we first plotted the RFU vs. temperature in no template controls (NTCs) used in the real-time PCR reactions (Fig. 1A). The NTC samples contain all reactants except for the template gDNA. The RFU of free fluorophore in these reactions exhibited a temperature-dependent linear decay in fluorescence across the entire temperature range tested (Fig. 1A). These results validate extrapolating the post-melt region to estimate background fluorescence from the unbound fluorophore to the earlier temperature points (see below).

**Fig. 1.**
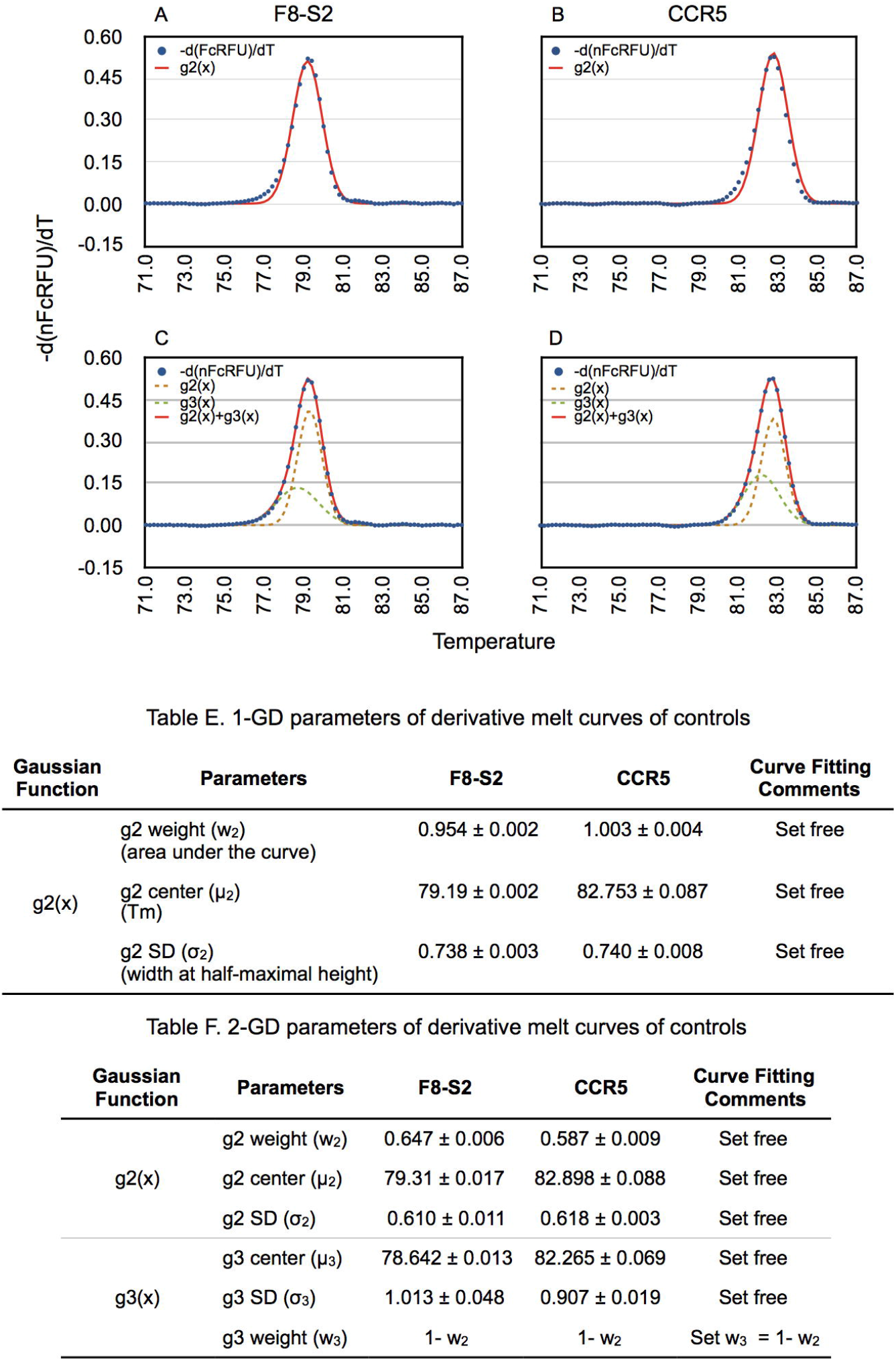
Temperature-dependent quenching of fluorescence of free and dsDNA-bound fluorophore and its correction. (A) Plot of first-order polynomial curve fit of raw RFU vs. temperature in no template controls (NTC). The equation shown in the plot is the mean ± SD of six different sample slopes and constants. (B) The unprocessed high-resolution melting profile (blue trace) and the extrapolation from first-order polynomial curve fitting of the post-melt curve region (red dashed line) from an amplicon of an unedited target site. (C) High-resolution melting profile of background subtracted RFU (BcRFU, blue trace) and that of ‘unquenched’ or fluorescence-compensated BcRFU (FcRFU, green trace) from an unedited target site. The red dashed line shows extrapolation of pre-melt region from first-order polynomial curve fitting of BcRFU and depicts the predicted BcRFU in the absence of DNA melting. D) Comparison of first-order polynomial curve fitting of post-melt and pre-melt portions of melting curves. Normalized data were used to enable plotting of the two sets of data.

For correction of background fluorescence for each melt curve, we carried out first-order polynomial curve fitting of the post-melt region of each melt curve data and then extrapolated the background RFU values for earlier temperature data points (red dashed line in Fig. 1B). We then subtracted the background RFUs corresponding to each temperature point to obtain the background subtracted RFU or BcRFU as described in Materials and Methods (Equation 2). The *BcRFU(x)* melt curve is shown in Fig. 1C (blue trace). The post-melt region of background subtracted-curve was nearly horizontal with an RFU close to zero indicating that the background fluorescence from free or unbound fluorophore was correctly computed and removed by this method.

### Correction of RFU for temperature-dependent quenching of dsDNA-bound fluorophore

To correct for quenching of fluorescence of dsDNA-bound fluorophore of background subtracted melt curve data (*BcRFU(x)*), we carried out a regression analysis of the pre melt region of mock-transfected samples and extrapolated the RFUs across the range of temperatures (red dashed line in Fig. 1C) (Equation 3). We obtained the efficiency of detection of dsDNA-bound fluorophore by normalizing *F*_prem_*(x)* to the estimated RFU at the starting temperature (*T*_low_ or 71°C) (Equation 4). The efficiency at each measured temperature was then determined for multiple mock samples (Fig. 1D). Measured efficiencies were nearly identical, diverging slightly at the higher temperatures, despite determination across experiments conducted on different days, and with different samples. The BcRFU of mock and test samples were divided by the average fluorescence efficiency at each measured temperature to obtain fluorescence corrected *BcRFU(x)* or *FcRFU(x)* ( Fig. 1C, green tracing) (Equation 5). The pre-melt region was now rendered horizontal and did not exhibit the temperature-dependent quenching profile of uncorrected melting curves. For the F8-S2 target amplicon melt curve fitting with a first order polynomial proved sufficient; for the CCR5 target amplicon melt curve, a second-order polynomial was required (see below).

We next wished to directly compare the temperature-dependent quenching effect on bound fluorophore vs. free fluorophore. To enable this comparison, we normalized the extrapolated background RFUs (determined from individual post-melt curve data of mocks) and plotted these along with the normalized bound-fluorophore efficiency (Fig. 1D). As anticipated from the NTC data shown in Fig. 1A, the slope of the free fluorophore (-0.002) was much more shallow than that of the bound fluorophore (-0.04). Thus, temperature-dependent fluorescence quenching of dsDNA-bound fluorophore is more pronounced and significant than that of the unbound or free fluorophore.

### Rationale for Gaussian modeling of first derivate melt curves

After high-resolution melt curve data were corrected for temperature-dependent quenching of unbound and dsDNA-bound fluorophore, curves were normalized and then numerically differentiated (Materials and Methods, Equations 6 and 7, respectively). When plotted, the processed data showed that both the pre-melt and post-melt regions were squarely placed on the zero baseline as expected (Fig. 2). The resulting peak of the first-derivate melt curve data resembled a “bell” curve. Bell-shaped density distribution curves can result from Cauchy-Lorentz, Student’s-t, Logistic or Gaussian distributions [19]. The Cauchy-Lorentz density distribution has longer tails, while the Student’s-t and Logistic density distributions exhibit heavier tails (kurtosis). The Gaussian distribution therefore seemed more suitable for empirical modeling of first-derivative melt curves. A preliminary curve fitting analysis using the Cauchy-Lorentz distribution function showed lower fit scores than the Gaussian distribution function.

**Fig. 2.**
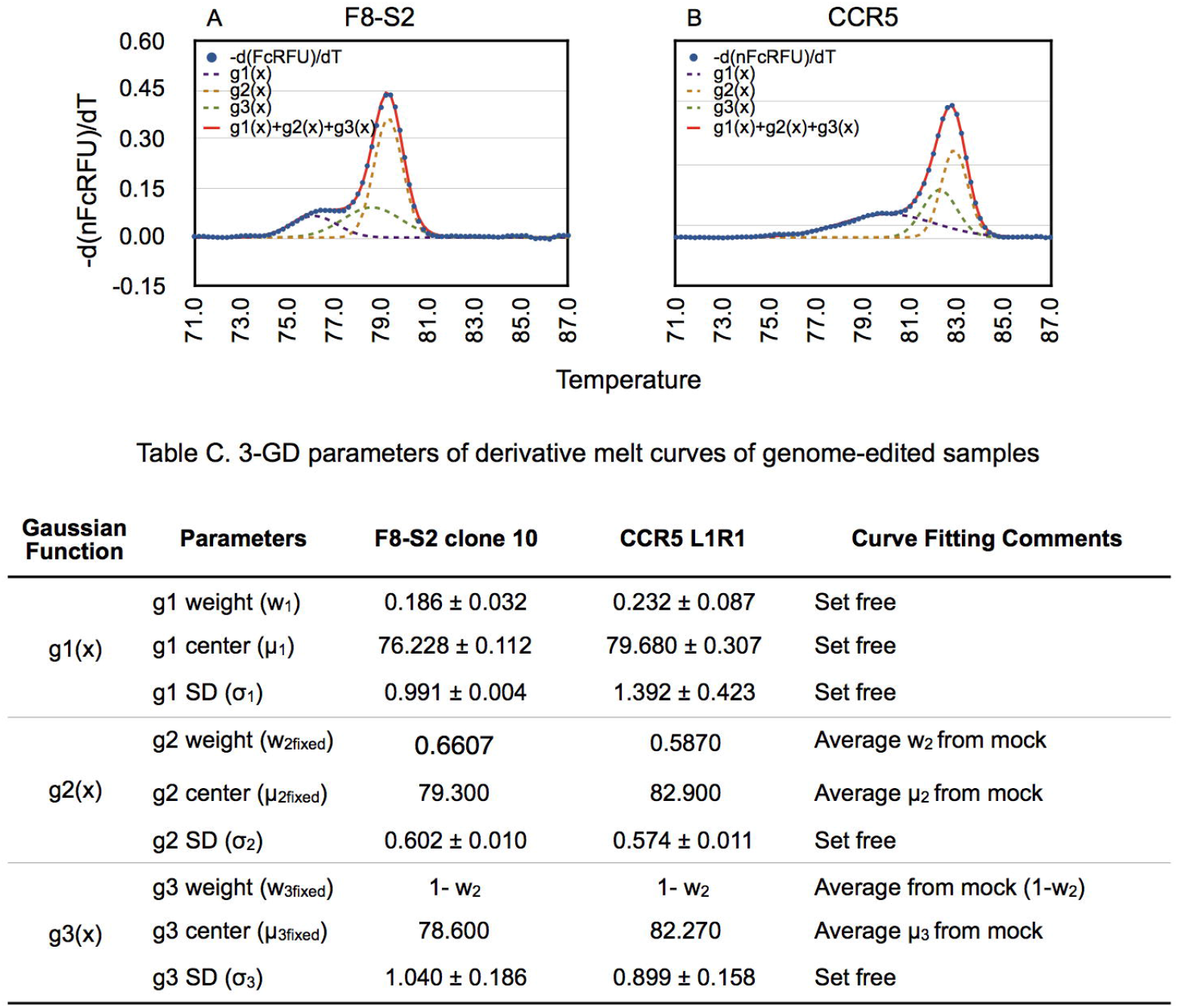
GD of first derivative of high-resolution melt curves of amplicons from gDNA of unmodified target sites. gDNA from mock-transfected HEK293T cells (Mocks) were PCR amplified using primer pairs targeting F8-S2 or CCR5 loci to obtain high resolution melt curve data as described in Materials and Methods. The normalized and fluorescence corrected melt curve data (nFcRFU) from F8-S2 (A and C) and CCR5 (B and D) target sites were numerically differentiated as described in Materials and Methods (Equation 7). 1-GD (A and B) and 2-GD curve fitting of derivative melt curves were done using CurveExpert Professional using Equation 9 and Equation 10, respectively. The first derivative (y-axis: -d(nFcRFU)/dT) was plotted against temperature (x-axis) and is shown as blue dots. The 1-GD curve fit to the first derivative data is shown as a red trace in A and B. The individual Gaussians of 2-GD curve fit are shown as brown (g3(x)) or green dashed lines and their sum (g2(x) + g3(x)) is depicted as a solid red line in C and D. Table E shows the Gaussian parameters determined from 1-GD curve fitting of A and B, while Table F shows the parameters identified by 2-GD curve fitting of C and D using the CurveExpert Professional software.

### Two-Gaussian decomposition is superior to one-Gaussian modeling of derivative melt curves of unmodified target sites

We first determined the parameters of the Gaussian components of first derivatives of *nFcRFU(x)* of unmodified or control samples (mocks) by curve fitting using the commercial software CurveExpert Professional(Materials and Methods). Gaussian curve fitting requires the user to input initial guesses for three of the parameters of a Gaussian function: curve weight (w), curve center (µ), and width at half-maximal height (σ) or standard deviation (SD). After multiple converging iterations using systematic changes to the parameters of the model, the software finds parameters with the fitting accuracy required or the maximum number of iterations is reached. The curve fitting output consists of the curve-fitted weight (‘w’ or area under the curve), curve center (µ) and the SD (σ). The better the curve fit, the closer the weight or area under the curve approaches 1 for derivatives of normalized melt curves.

We wished to use the simplest possible mathematical model for measuring the proportion of mutant population in the amplicon of the target region. This would consist of one Gaussian component for describing first derivative of nFcRFU of unmodified mocks and another Gaussian for the mutant population. The first derivative melting curves (-*d(nFcRFU(x))/dx*) from unmodified F8-S2 and CCR5 loci (Fig. 2A and 2B) were curve fitted using a single-Gaussian function, *g2(x)* (Materials and Methods, Equation 9). We refer to this as single-Gaussian decomposition (1-GD). Modeling the first derivative of the F8-S2 target site showed the area under the curve had a weight (*w_2_*) of 0.9537 ± 0.0021. The deviation of the fitted curve from the actual melt curve was clearly visible over the pre-melt to melt transition region where the Tm of the amplicons with deletion mutations is situated (Fig. 2A). 1-GD curve fitting for the CCR5 target was similar to that of F8-S2 target but with only a slight divergence from the actual derivative melt curve (Fig. 2B). Consistent with this the area under the curve was 1.003 ± 0.0039 (from four independent replicates). As in the case of the F8-S2 target site, we saw a small divergence in the early melting region Fig. 2B (*g2(x)* vs. -*d(nFcRFU)/dT*).

Since the mutant molecules contribute to the melt profile in the early melt region, it was necessary to ensure a more accurate curve fitting over this region than provided by a single Gaussian component. To this end, we tested modeling of derivative melt curves of unmodified controls as a sum of two Gaussian functions, *g2(x)* + *g3(x)*, (Materials and Methods, Equation 10). As for the 1-GD curve fitting, we provided initial best guesses for the five parameters (three for first Gaussian component and two for the second Gaussian component). For the g3(x) Gaussian we suggested initial guesses for the mean (*µ3*) over the pre-melt/melt transition region. We stipulated that the sum of weights for *w*_2_ and *w*_3_ should equal one and set free *w_2_* (and thereby, *w3= 1-w2*). The results of this curve fitting experiment are shown in Fig. 2C and 2D for the F8 and CCR5 loci, respectively. Unlike 1-GD curve fitting, the sum of two Gaussian curve fitting (Fig. 2, g2(x) + g3(x) indicated by a red trace vs. -d(*nFcRFU*)/dT indicated by blue dots) recreated the derivative melt curve nearly perfectly. When we compared CurveExpert Professional scores (see Materials and Methods, Comparing two curve fitting models), the two-Gaussian decomposition (2-GD) model outscored the 1-GD model for both F8 and CCR5 mock samples (Table 1). This difference, although slight, was statistically significant (paired Student’s-t test, p = 0.0000). The AICc values were lower for the 2-GD model indicating that it had a better fit. The relative likelihood calculations from the AICc values of both 1- and 2-GD models, also showed that 2-GD model was better (Table 1).

**Table 1.** 2-GD model shows better fit than 1-GD for derivative melt curve data of mocks

| Targeting<br>Plasmid | Selection | Sample | 1-GD<br>score | 2-GD score | 1-GD AICc | 2-GD AICc | ΔAICc | Relative<br>likelihood |
| --- | --- | --- | --- | --- | --- | --- | --- | --- |
| pBackbone | No | F8-S2 Mock 1 | 981 | 995 | -625 | -785 | 160 | 0.0000 |
| pBackbone | No | F8-S2 Mock 2 | 982 | 995 | -629 | -795 | 166 | 0.0000 |
| pDonor | No | F8-S2 Mock1 | 980 | 995 | -620 | -786 | 166 | 0.0000 |
| pDonor | No | F8-S2 Mock 2 | 980 | 996 | -618 | -812 | 194 | 0.0000 |
| pBackbone | Yes | F8-S2 Mock 1 | 978 | 996 | -615 | -813 | 198 | 0.0000 |
| pBackbone | Yes | F8-S2 Mock 2 | 979 | 996 | -618 | -803 | 185 | 0.0000 |
| pDonor | Yes | F8-S2 Mock 1 | 981 | 995 | -629 | -791 | 161 | 0.0000 |
| pDonor | Yes | F8-S2 Mock 2 | 981 | 996 | -625 | -806 | 181 | 0.0000 |
| None | No | CCR5 Mock 1 | 984 | 997 | -732 | -979 | 247 | 0.0000 |
| None | No | CCR5 Mock 2 | 985 | 998 | -748 | -969 | 221 | 0.0000 |
| None | No | CCR5 Mock 3 | 984 | 998 | -729 | -1057 | 328 | 0.0000 |
| None | No | CCR5 Mock 4 | 983 | 997 | -721 | -990 | 268 | 0.0000 |

First-derivative melt curves from unmodified F8-S2 and CCR5 target sites provided distinct Gaussian parameters from curve fitting as expected from their differing amplicon sizes, sequences and differing Tms. Thus, they exhibited distinct centers or means for both 1-GD (*µ_2_* of 79.19 ± 0.002 vs. 82.753 ± 0.087) (Table E in Fig. 2) and 2-GD fitting (*µ_2_* of 79.31 ± 0.017 vs. 82.898 ± 0.088 and *µ_3_* of 78.642 ± 0.013 vs. 82.265 ± 0.069 for F8-S2 and CCR5, respectively) (Table F in Fig. 2). Likewise, they showed distinct differences in the contribution of weights: *w_2_* of 0.954 ± 0.002 vs. 1.003 ± 0.004 in 1-GD fitting; and *w_2_* of 0.647 ± 0.006 vs. 0.587 ± 0.009 for F8-S2 and CCR5, respectively in 2-GD fitting. These results highlight the requirement for determining Gaussian parameter values for each target site from amplicons obtained from corresponding control or unmodified samples.

### Estimating percentage of mutants by GD of derivative melt curves from genome-edited samples

Comparing derivative melt curves of unmodified and genome-edited samples shows a distinct mutant molecules’ peak with a lower melting temperature (Fig. 2 vs. Fig. 3). *We hypothesized that upon decomposition of the melting profile into its Gaussian components, the area under the mutant peak would correspond to the proportion of mutant molecules in the PCR product*. The Gaussian function representing the mutant population was designated *g1(x)* in Equations 11 and 12 (Materials and Methods).

**Fig. 3.**
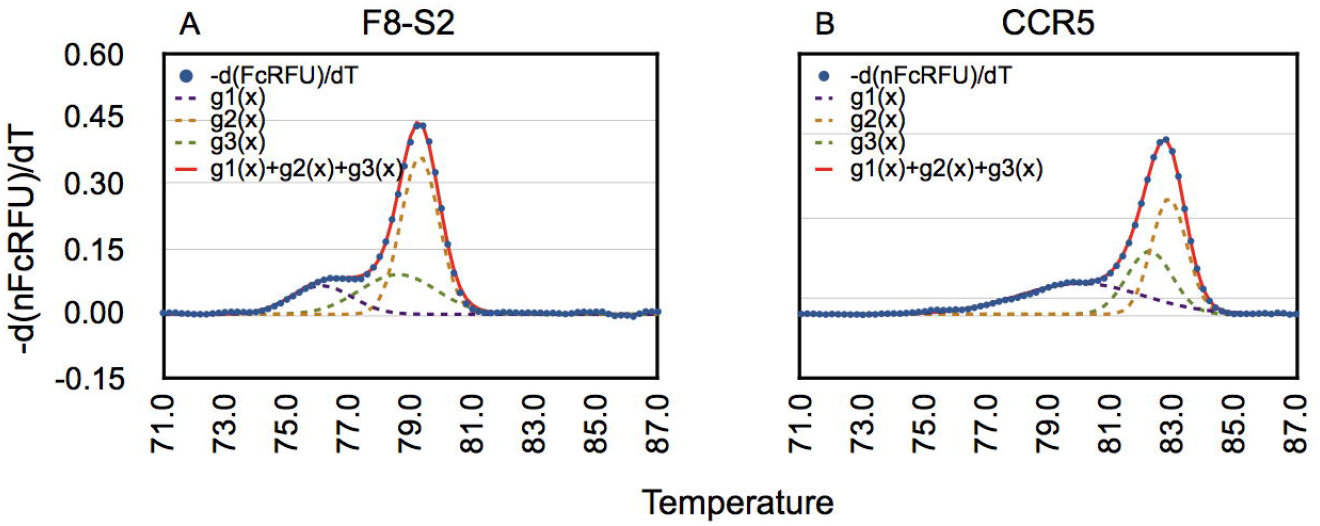
3-GD of first derivative of high-resolution melt curves for estimation of mutant percentage in genome-edited samples. gDNA was isolated from HEK293T cells transfected with F8-S2 targeting RGENs or CCR5 targeting TALENs and PCR amplified using corresponding primer pairs to obtain high resolution melt curve data (Materials and Methods). 3-GD curve fitting was done on first derivative melt curves using CurveExpert Professional and Equation 12 as described in Materials and Methods. The individual Gaussians-g1(x) (purple dashed line), g2(x) (brown dashed line) and g3(x) (green dashed line) and their sum-g1(x)+ g2(x) + g3(x) (red solid line) were overlaid over the first derivative melt curve (blue dots). GD of F8-S2 is shown in A and of CCR5 in B. Table C shows the parameters (weights, centers and SDs) of 3-GD. The parameters that were fixed from GD of mocks and those that were set free during 3-GD of edited samples are shown in the Comments column. The g1 weight (w_1_) represents the mutation frequencies in the amplicons of genome-edited F8-S2 and CCR5 target sites, respectively.

Since the better curve fitting of unmodified controls was obtained by using sum of two Gaussian functions, we modeled derivative melt curves of test samples as a sum of three Gaussian functions, *g1(x)* + *g2(x)* + *g3(x)* (Materials and Methods, Equation 12). The parameters obtained from 2-GD of derivative melt curves of unmodified controls from F8-S2 and CCR5 (means and weights) were then used to decompose corresponding test or genome-edited samples. The different Gaussian components, g1(x), g2(x) and g3(x), and their sum g1(x)+ g2(x) + g3(x) are shown in Fig. 3. The predicted curve of the sum of the three Gaussians was a near-perfect fit to the original derivative melt curve from test samples (Fig. 3, g1(x) + g2(x) + g3(x), indicated by a red tracing vs. *-d(nFcRFU)/dT* (Fig. 3, blue dots). The area under the g1 curve, *w*1, of three-Gaussian decomposition (3-GD) was deemed to represent the mutant population. The percentage of mutant population estimated in amplicons of genome-edited F8-S2 and CCR5 target sites by 3-GD, shown in Table C in Fig. 3, was 18.6 ± 3.2% vs. 23.2 ± 8.7%, respectively. These results demonstrate that first derivative melt curves from genetically altered sites can be modeled successfully as a sum of three Gaussian functions.

Since the 1-GD of unedited samples was below the data points in the pre-melt to melt transition region (Fig. 2), we hypothesized that 2-GD of genome-edited samples would over estimate the mutant frequency. The results of these comparisons are shown in Fig. 4. 2-GD modeling estimated significantly higher mutant frequency than 3-GD modeling of edited samples (Fig. 4A and 4B) as predicted.

**Fig. 4.**
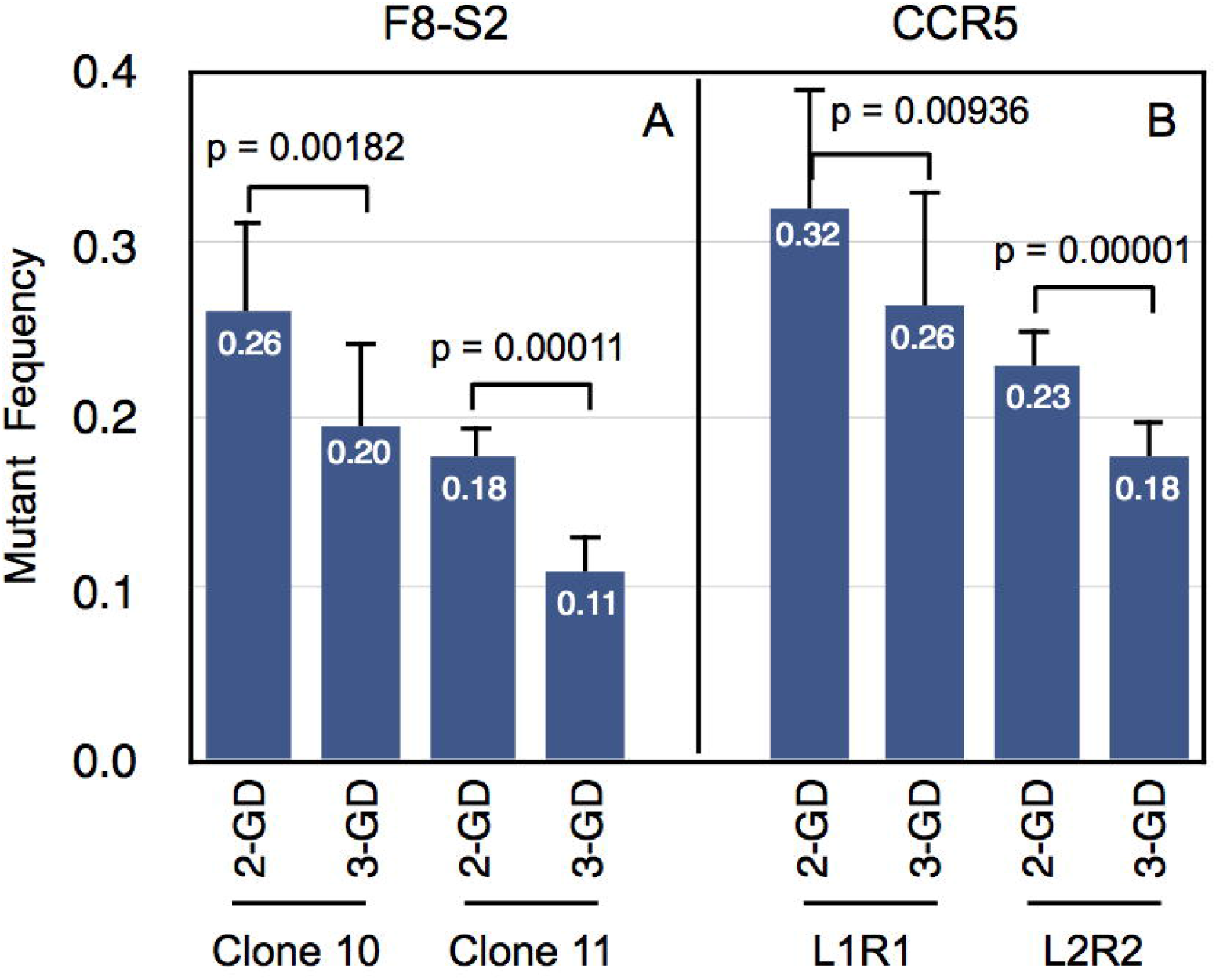
Comparison of mutant percentage estimation by 2- and 3-GD. First derivatives of high-resolution melt curves from genome-edited samples were curve fitted using 2- or 3-GD models as described in Materials and Methods (Equation 11 and Equation 12, respectively). The mutant percentages estimated from curve fitting are shown along the y-axis for F8-S2 (A) and CCR5 (B). Two molecular clones (10 and 11) of dgRNAs targeting F8-S2 site and two pairs of TALENs (L1R1 and L2R2) targeting CCR5 site were tested. The mutant percentages were compared using Student’s t-test (two-tailed). The p-values of the pair-wise comparisons of 2-GD and 3-GD are shown above the bars.

Better curve fitting of 3-GD over 2-GD modeling was also revealed by the CurveExpert Professional scores (Table 2). These differences were statistically significant (paired Student’s t-test, p = 0.00001). The AICc values were lower, indicating a better fit, for the 3-GD model. Relative likelihood determinations from AICc values also revealed that the 3-GD model was better. These results demonstrated that the 3-GD modeling was the appropriate choice for GD of first derivative melt curves of amplicons of genome-edited target sites.

**Table 2.** 3-GD model achieves better fit than 2-GD for derivative melt curve data of genome-edited samples

| F8-S2 site |  |  |  |  |  | CCR5 site |  |  |  |  |  |
| --- | --- | --- | --- | --- | --- | --- | --- | --- | --- | --- | --- |
| 2-GD<br>Score | 3-GD<br>Score | 2-GD<br>AICc | 3-GD<br>AICc | ΔAICc | Relative<br>likelihood | 2-GD<br>Score | 3-GD<br>Score | 2-GD<br>AICc | 3-GD<br>AICc | ΔAICc | Relative<br>likelihood |
| 979 | 992 | -780 | -911 | 131 | 0.0000 | 990 | 997 | -837 | -988 | 151 | 0.0000 |
| 990 | 996 | -879 | -1005 | 126 | 0.0000 | 985 | 995 | -776 | -907 | 130 | 0.0000 |
| 992 | 996 | -923 | -1008 | 85 | 0.0000 | 978 | 993 | -733 | -870 | 137 | 0.0000 |
| 983 | 994 | -797 | -949 | 153 | 0.0000 | 968 | 990 | -690 | -830 | 140 | 0.0000 |
| 990 | 995 | -871 | -978 | 108 | 0.0000 | 978 | 993 | -728 | -871 | 143 | 0.0000 |
| 992 | 996 | -888 | -1000 | 112 | 0.0000 | 973 | 992 | -704 | -857 | 154 | 0.0000 |
| 991 | 996 | -869 | -980 | 111 | 0.0000 | 972 | 992 | -703 | -851 | 148 | 0.0000 |
| ND | ND |  |  | ND | ND | 968 | 991 | -681 | -834 | 152 | 0.0000 |

### Comparison of GD method to prior approaches for measuring efficiency of genome editing

We next carried out 3-GD of high resolution melt curves of samples previously characterized by NGS and by an alternative approach to measure mutant population based on difference curve areas (DCAs) of normalized high-resolution melt curve profiles. These samples exhibited a wide range of mutant percentages that were influenced by puromycin drug selection and the use a donor template containing plasmid (pDonor-F8) or its corresponding control plasmid (pBackbone) [12]. There were four categories of samples: (1) pBackbone/Unselected, (2) pDonor/Unselected, (3) pBackbone/Selected, and (4) pDonor/Selected. These four categories showed progressively increasing percentages of mutations in the earlier study [12]. Two different clones of RGENs targeting the F8-S2 site, clone 10 and clone 11, were tested. Clone 10 had previously exhibited higher efficiencies than clone 11.

Results of curve fitting of derivative melt curves of mocks using 2-GD and of genome-edited samples by 3-GD are shown for all the replicate samples in Fig. 5A. In all instances, GD was able to accurate model the derivative melt curves including the mutant molecules’ peak. The area under this peak, w_1_, is shown as percentage within the plots. RGEN F8-S2 clone 10 edited samples showed higher percentages of mutants than clone 11. Drug-selected samples exhibited higher mutant frequencies than corresponding unselected samples and samples that received pDonor-F8 template (to effect homologous recombination) exhibited higher mutant frequencies than corresponding samples that received the control pBackbone plasmid.

**Fig. 5.**
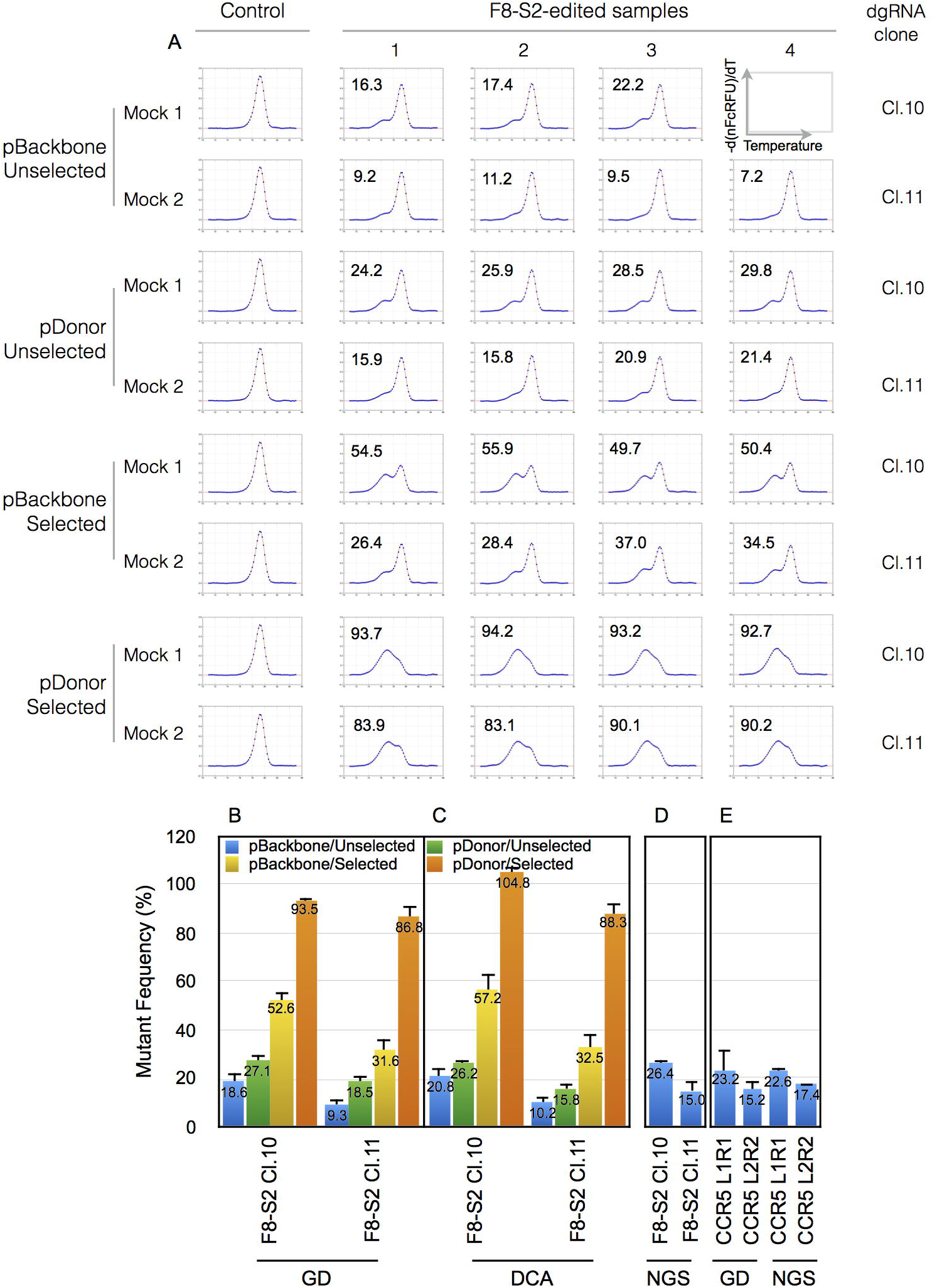
Mutant frequency determination by 3-GD and comparison to difference curve areas (DCAs) and next generation sequencing (NGS) data. HEK293T cells were transfected with F8-S2 targeting dgDNA clone 10 (F8-S2 Cl.10) or clone 11 (F8-S2 Cl.11) together with a dCas9-FokI construct. The cells were also cotransfected with either pBackbone or pDonor-F8 targeting plasmids (Materials and Methods). Following transfection, gDNAs were isolated from unselected cells or cells selected with puromycin and used for amplification by PCR using appropriate primer pairs targeting F8-S2 loci to obtain high-resolution melt curve data. (A) Mutant percentage estimations by 3-GD for the four different categories of samples from unedited and edited F8-S2 site are identified on the left. The derivative melt curves are shown as blue dots and the fitted curves from GD as red traces. Four PCR replicates were analyzed for each clone with one exception (F8-S2 clone 10, pBackbone/Unselected) for which only three replicates were tested. The mutant frequency (percentage) estimated from the area of the mutant peak (*w1* parameter from *g1(x)*), of 3-GD) for each replicate is shown within the plot. (B-D) The average mutant frequency determined by GD for the different categories in A were compared to mutant frequencies determined by difference curve areas (DCA) (C) and to mutant frequency determination from next generation sequencing (NGS). NGS was only done on unselected samples. (E) Mutant frequency estimation from GD of high resolution melt curve data from gDNA of HEK293T cells transfected with TALENs (two independent pairs of molecular clones L1R1, L2R2) targeting CCR5 locus. CCR5 edited samples were also analyzed by NGS. Error bar = 1 SD.

Direct comparison of the results with mutant frequency determination using DCA is shown in Fig. 5B-C. Consistent with our previous observations, the percentage of mutants estimated by both methods were within 3% of each other for both selected and unselected samples (pBackbone or pDonor). There were two exceptions where the differences were 4.6% and 11.3%, respectively, with GD providing lower estimates. Possible explanations for this discrepancy are provided in Discussion. The NGS of unselected samples treated with pBackbone showed a similar trend as the above two methods (Fig. 5D) with clone 10 again showing higher efficiency of target site modification than clone 11. NGS generally provided higher estimations of mutant frequencies than GD or DCA methods due to the inclusion of insertion mutations in the calculations.

We used GD to also estimate the proportion of mutants in amplicons of samples edited at the CCR5 locus. Here too, the results of GD and NGS showed similar trends (Fig. 5D). These results *in toto* demonstrate that curve fitting of first derivative of high-resolution melt curves is comparable to other methods used previously for estimating the proportion of mutants in amplicons of genome-edited target sites. The results also indicate that one could estimate mutant frequency percentages by GD for target sites for which there is no ready availability of a 100% mutant population to generate calibration curves for the DCA method (in this case genome-edited CCR5 target site).

### The size of the PCR product does not affect estimation of percentage of mutants by GD from the same target locus despite exhibiting distinct Gaussian parameters

We next wished to test if the size of the amplicon affected the estimation of percentage of mutants. To this end, we amplified unmodified or genome-edited CCR5 target sites using two sets of primers. The same antisense primer (SK145) was used for both PCR amplifications but one of the sense primers (SK214) was situated further upstream of primer SK144 so that the resulting amplicon sizes were 140 and 107 bp, respectively. GD of high-resolution melt curves of both sizes of amplicons was done as above. Results are shown in Fig. 6. The larger amplicon exhibited higher means (*µ_1_*, *µ_2_* and *µ_3_*) for the three Gaussian functions than the smaller one, as expected, and also showed distinguishable SDs (Table 3). The percentages of mutants estimated from the larger or smaller PCR product sizes determined by GD were 29.8 ± 1.1 % vs. 28.9 ± 8.6 %, respectively. The values were not statistically significant (Student’s t-test, p≥0.05). These results suggest that small differences in amplicon sizes (less than 50 bp) do not affect the estimation of genome-editing efficiency by GD.

**Fig. 6.**
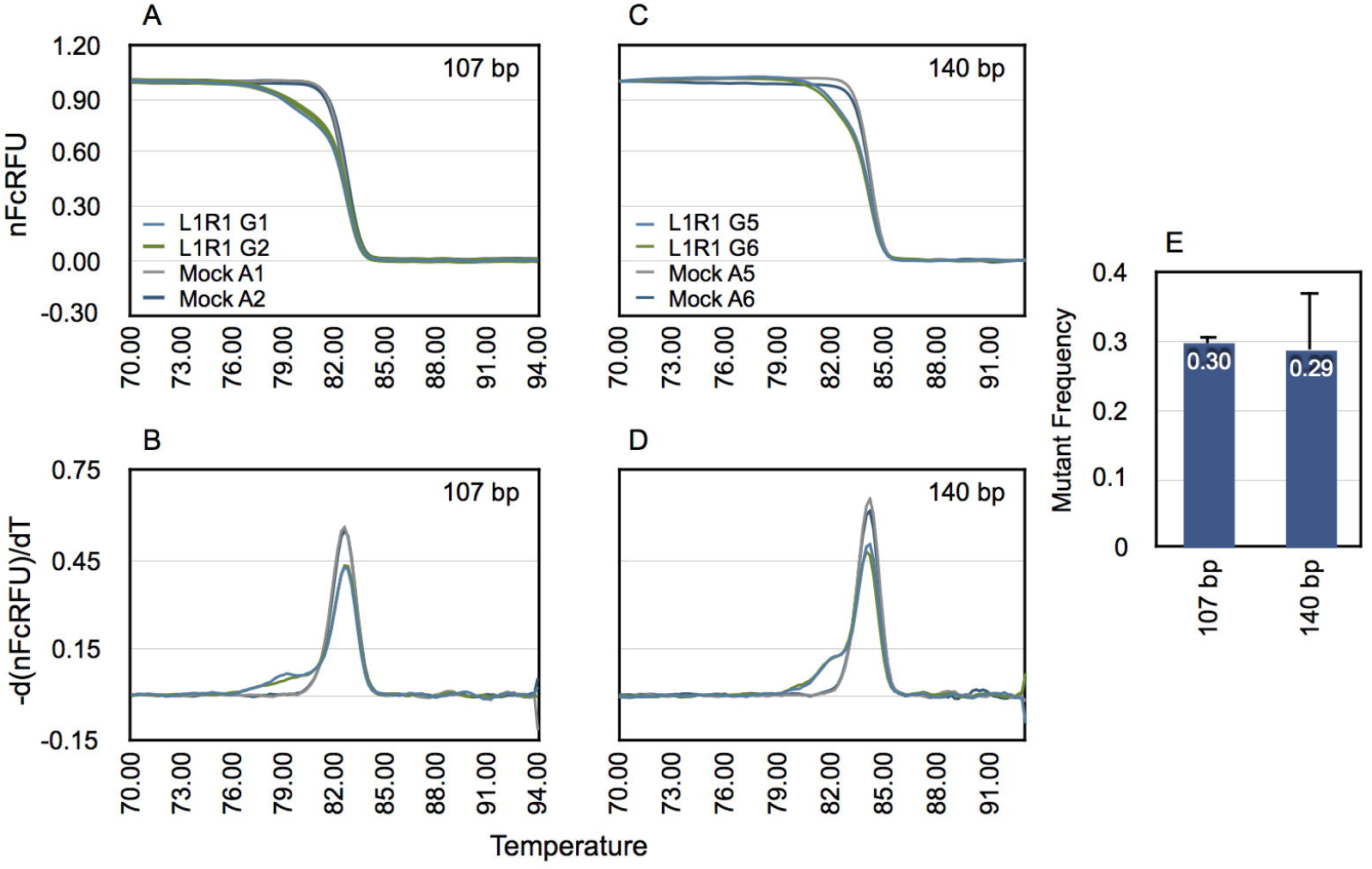
Size of PCR product does not affect determination of mutant percentage by GD. The CCR5 target site in gDNA of unmodified or genome-edited cells were amplified using two pairs of primers designed to produce two distinct sizes of product (107 bp and 140 bp, respectively). The amplicons were subjected to high-resolution melting and then processed to correct for temperature-dependent quenching of fluorescence of free and dsDNA-bound fluorophore. The resulting melt curves of genome-edited (for clone pair L1R1) and unmodified controls (Mock) are shown (A & C). Corresponding first-derivatives of processed melt curves are shown in B and D. Replicates G1 and G2, A1 and A2 refer to gDNA samples amplified using primers that produce 107 bp amplicon, whereas G5 and G6, and A5 and A6 refer to gDNA samples amplified using primers that produce 140 bp amplicon. The derivative melt curves were decomposed using the 3-GD model to estimate the mutant frequency. The estimated mutant frequencies for both sizes of amplicons are shown in (E). Error bar = 1 SD.

**Table 3.** Parameters determined by 3-GD of two different size amplicons from the CCR5-edited target site

| Gaussian Parameters | 107 bp PCR product | 140 bp PCR product |
| --- | --- | --- |
| $W_1$ (%) | $29.8 \pm 1.1$ | $28.9 \pm 8.6$ |
| $\mu_1$ | $79.8 \pm 0.23$ | $82.1 \pm 0.19$ |
| $\sigma_1$ | $1.88 \pm 0.20$ | $1.07 \pm 0.26$ |
| $w_2$ | $0.49 \pm 0.01$ | $0.48 \pm 0.06$ |
| $\mu_2$ | 82.8 | 84.2 |
| $\sigma_2$ | $0.57 \pm 0.01$ | $0.51 \pm 0.01$ |
| $w_3$ | $0.21 \pm 0.00$ | $0.24 \pm 0.03$ |
| $\mu_3$ | 82.1 | 83.6 |
| $\sigma_3$ | $0.77 \pm 0.05$ | $0.65 \pm 0.12$ |

## Discussion

Here we outline a method for estimating the efficiency of genome-editing reagents by GD of high-resolution melt curve data. An initial pre-processing of the raw melt curve data was required to correct for the quenching effect of temperature on measurement of fluorescence as a prelude to GD for estimating the genome-editing efficiency. Our approach consisted of two separate steps for correcting melting curves for temperature-dependent quenching of fluorophore. The initial step of cleaning the data involved removing the background fluorescence emanating from the free or unbound fluorophore. Two methods have been used for this purpose. The first is to use an arbitrary cutoff point in the post-melt region of the raw melt curve and subtract this value from all upstream RFUs. We found that this method sometimes resulted in a small but narrow tail in the postmelt region of the curve before it hit the baseline. This discrepancy could affect curve fitting of the first derivate of the processed melt curve. The tail also hinted at a temperature-dependent quenching of the free fluorophore. We confirmed this quenching from linear regression analysis of no template controls used in PCR across the entire range of melting (Fig. 1). The computed background RFU from linear regression of the post-melt region of individual melt curves was used to effectively subtract the effect of free fluorophore on the melt curve.

The second step to processing the melt curve involved correcting for temperature-dependent quenching of the dsDNA-bound fluorophore evidenced in the pre-melt region. As for the post-melt region, regression analysis of the pre-melt region can be used to determine the efficiency of fluorescence of the dsDNA-bound fluorophore at any temperature point along the melt curve profile. While detection efficiency can be computed for individual melt curve profiles, we found that the temperature range of the pre-melt region could be much shorter for some genome-edited samples due to the expected lower Tms for deletion mutations. For example, the pre-melt regions were only nominally present for drug-selected samples that had a very high proportion of mutant molecules in the amplicon (Fig. 5). In this case the mutant population constituted more than 90% of the PCR product.

We found that for a given target, and pair of primers, the efficiency of detection of dsDNA-bound fluorophore could be computed accurately and solely from unmodified or mock-transfected samples. These efficiencies could not be distinguished from those estimated from the individual test samples where sufficient pre-melt region was present (Fig. 1D). We therefore chose to determine bound fluorophore detection efficiency from replicates of mock-transfected samples and averaging them. Correction for the quenching of fluorescence of dsDNA-bound fluorophore could be simply achieved by dividing the *BcRFU(x)* by the detection efficiency, *E(x)* (Materials and Methods, Equation 4). This process effectively eliminated the downward slope of the pre-melt region (Fig. 1C).

The temperature-dependent decay of fluorescence of dsDNA-bound fluorophore could be modeled using either a first-or second-order polynomial function. For CCR5 samples, the pre-melt region, following a correction using a first-order polynomial, showed a gentle upward trajectory (saddleback pre-melt region) indicating that the RFU was not compensated appropriately. Estimating the dsDNA-bound fluorophore efficiency using a second-order polynomial curve fitting of the pre-melt region eliminated this artifact. From this one can surmise that the fluorescence decay of dsDNA-bound fluorophore at higher temperatures is better modeled with a second-order polynomial.

Correction for temperature-dependent quenching of fluorophores has been described previously. Watras et al., found that fluorescence of chromophoric dissolved organic matter (CDOM) decreased as ambient water temperature increased [20]. They suggested compensating for the quenching using the equation:

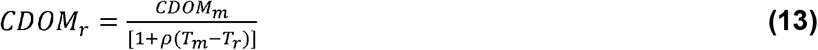

where t = temperature (°C), r= reference and m = measured values, the coefficient, ρ, is the quotient of slope divided by the intercept. The actual coefficient value, ρ, was found to be instrument-dependent. A similar approach was recommended by Ryder et al [21,22].

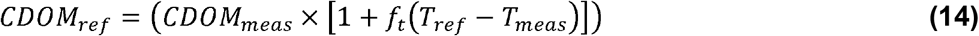

where *f*_t_ is the temperature correction coefficient, *ref* and *meas* refer to reference and measured temperatures. The two formulae for calculating fluorescence compensation were shown to be mathematically identical [23]. This correction method is comparable to our approach. Our initial attempts at correction for the quenching effect was to determine the slope of pre-melt region and use it in place of the coefficient, ρ, in Equation 13. This was combined with a simple baseline cut off for correction of melt curve data. We, however, prefer first-order polynomial curve fit to determine and subtract the background from individual melting curves, and then correct for the quenching effect of temperature on dsDNA-bound fluorophore by dividing with the efficiency of detection of dsDNA determined from unmodified controls. Both approaches should provide comparable results for subsequent curve fitting after numerical differentiation. Our approach eliminates the requirement for slope determination of the pre-melt region for each of the test samples easing computation.

Palais and Wittwer described two methods for background correction [24]. 1) A baseline method:

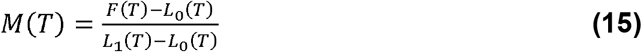

where, *M(T)* is the corrected melt curve, *F(T)* is the experimentally obtained melt curve, and *L*1*(T)* and *L*_0_*(T)* refer to linear equations describing pre-melt and post-melt regions of the curve, respectively. Thus, *M(T)* corresponds to *FcRFU(x)*, *F(T)* to *RFU(x)*, *L*_1_*(T)* to *F*_prem_*(x)* and *L*_o_*(T)* to *B*_pom_*(x)* of this study.

2) They also described an exponential background subtraction model:

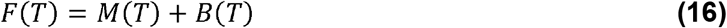

Where the background, *B(T)=Ce^a(T-T_L_)^ C* and *a* are determined as described in detail in their publication.

The exponential background correction is recommended by Palais and Wittwer for experiments involving multiple small amplicons and unlabeled probes, and also where the pre-or post-melt regions of melt curve exhibit a concavity. We evaluated the exponential background subtraction method to process the raw melting curve data for amplicons of F8-S2 and CCR5 loci in unedited mock samples. The results are shown in Figs. 7A and 7B and indicate that this correction method only partially compensated for the quenching observed in the pre-melt region. Since the mutant population encroaches on pre-melt region and extends into the melt transition portion, we abandoned this approach for preprocessing the high-resolution melt curves.

**Fig. 7.**
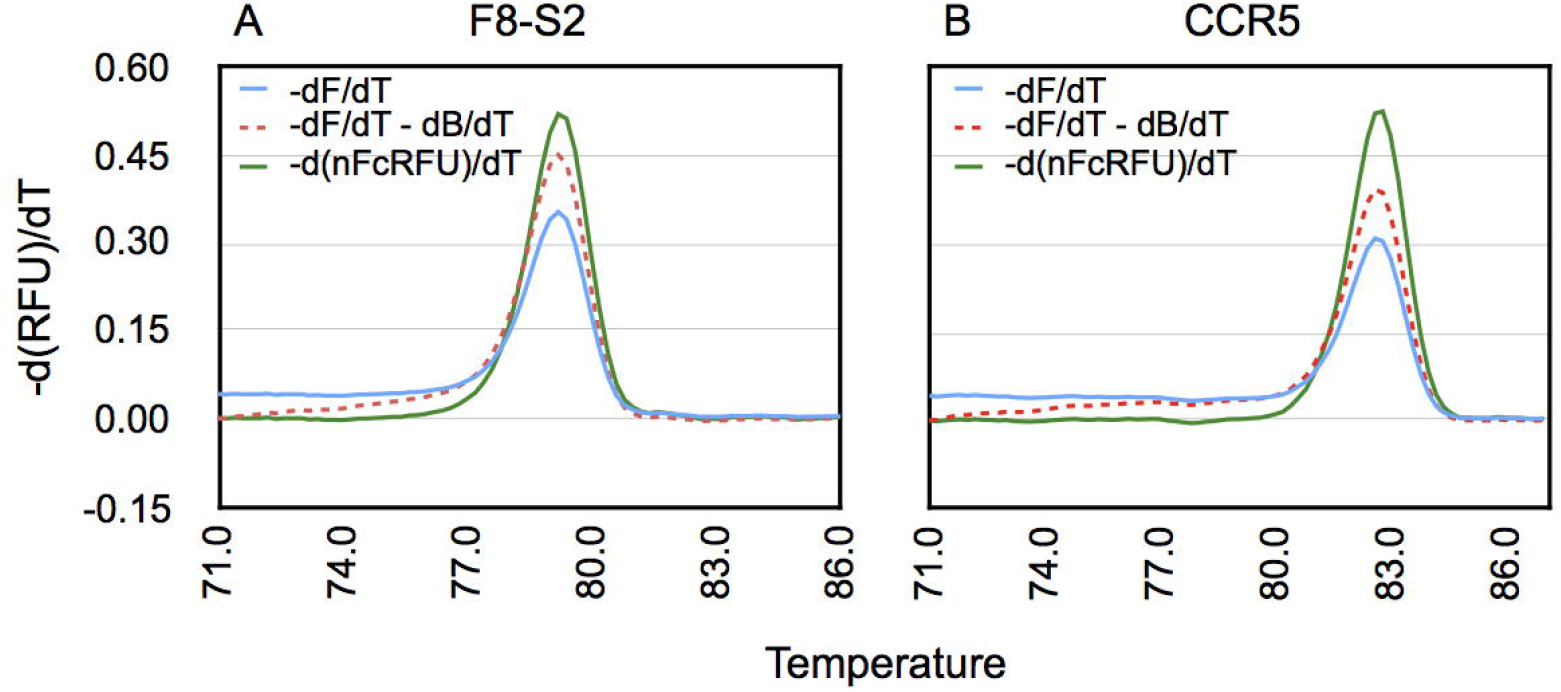
Comparison of different methods of processing melt curve data for background and fluorescence quenching correction. Melt curve data from amplicons of unmodified or control samples from F8-S2 (A) or CCR5 target loci (B) were either unprocessed (-dF/dT, blue trace) or corrected using exponential background subtraction method of Palais and Wittwer (24) (-dF/dT-dB/dT, red dashes) or the method described in this study (-d(nFcRFU)/dT, green trace).

Our method for preprocessing melt curve data is mathematically indistinguishable from the simpler baseline model of Palais and Wittwer (Equation 15). One difference between the Palais and Witter method and our method is that we first subtract background emanating from unbound fluorophore before correcting for efficiency of detection of dsDNA-bound fluorophore. The second difference is that we formulate the decrease in fluorescence of the pre-melt region not as a background problem but rather as an issue of detection efficiency. The third difference is that the quenching of dsDNA-bound fluorophore was modeled using either a first-or a second-order polynomial function depending on the particular target amplicon. The final difference is that we determined ds-DNA bound fluorophore detection efficiencies from control or mock samples and applied those to correct melt curves of genome-edited samples.

After preprocessing melt curve data, we used GD to successfully model first derivative melt curves. Cuellar and coworkers were amongst the earliest investigators to analyze high-resolution denaturation profiles of reassociated repetitive DNA sequences using a combination of higher derivative analysis and curve fitting [25]. They were able to distinguish “thermal classes” of repetitive DNA duplexes exhibiting different amounts of base pair mismatch in reassociated DNA. Reassociated *Escherichia coli* DNA exhibited a single thermal class while pea and mung bean re-associated DNAs showed five distinct thermal classes. These investigators obtained the first to fifth derivatives of the melting profiles by numerical differentiation followed by smoothing using nine-point running averages. For curve fitting of first derivative curves they used a software program called RESOLV. Their results showed that the number of peaks identified by RESOLV corresponded well with the fifth derivative of the melting profiles of reassociated mung bean or pea DNAs. While these investigators were able to use an empirical approach to identify multiple Gaussian components in reassociated DNA of legumes, they were unsure if the components corresponded to populations of distinct sequences.

Moore and Gray proposed a method dubbed derivative domain fitting for resolving a mixture of normal distributions in the presence of a contaminating background [26]. They proposed this model for analyzing flow cytometric data. A requirement for decomposition was that Gaussian peaks had to be separated by an SD greater than two. They mentioned difficulties in accurately modeling the background by their method. While their approach is an example of GD of data, their study is not directly comparable to ours.

Nellåker and coworkers proposed a *mixture model* to analyze of melting temperature data [27]. The premise of their model is that distinct Tm categories indicate presence of population of unique sequences. The “mixture model” allows calculating the proportions of amplicons contributing to the distinct Tm categories identified in the mixes. Nellåker and coworkers state that their *mixture model* actually denotes *mixture distributions* of statistical distributions that arise from sampling of mixed populations. They formulate the probability density function, g(x) as follows:

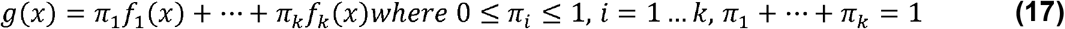

The parameters *π*_1_ … *π*_k_ are referred to as the mixing weights or proportions. They applied the mixture models to Tm data assuming it to consist of normally distributed components with each component having the same standard deviation. They used a Gaussian distribution function for their model. Thus, the function *g(x)* (Equation 17) was represented as:

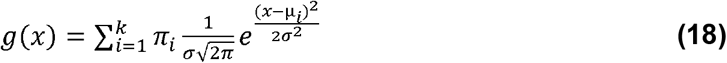

where, *x* refers to temperature, and *μ_i_* refers to Tm of individual components of the mixture

The sum of Gaussian functions that we used in this study (Materials and Methods, Equations 9 and 10) to curve fit the first derivative of processed melt curves, is similar to that of Nellåker and coworkers. However, Nellåker and coworkers used their Gaussian function for modeling Tm distributions of individual components of their mixture and did not apply it to derivative transformations of melt curves of mocks. Here, we *apply the sum of Gaussian functions to empirically reproduce the shape of the first derivative of high-resolution melt curves for both mocks (sum of two Gaussians) and genome-edited samples (sum of three Gaussians)*. A second difference is that we did not assume the SD was the same for the decomposed Gaussian components. They were designated as separate parameters for each Gaussian and set free during the modeling. However both Gaussian models sought to measure the proportion of particular component of the mixes, the only difference being, we designated the weight of the different components as *w*_1-3_ instead of π_i_. This also eliminated possible confusion between the weight coefficient and the mathematical constant π. In our case too, the sum of the weights of the Gaussian components of first derivative melt curves equaled one.

Mann et al., also used a Gaussian model to curve fit melt curve derivatives [28]. They were interested in automating the screening of first derivative melt curves following PCR to detect products with unusual or aberrant melt curves to rapidly eliminate those samples from further analyses. They used a different background correction method than those described above. Their approach provides a pure Gaussian after subtraction of a sigmoid shaped background fluorescence that does not retain the granularity of the derivative melt curve from genome-edited target sites. In our model, the shape of the derivative melt curve is critical for the precise quantitative decomposition into its Gaussian components.

There was good correspondence between the results obtained by GD and our earlier described method based on DCAs for estimating mutant amplicon frequency (Fig. 5). The DCA method was previously validated from NGS of the same amplicons. While the GD and DCA methods yielded comparable estimation of editing efficiencies, there were a few exceptions for amplicons consisting almost entirely of mutant species (Fig. 5A vs. 5B, pDonor/Selected samples). We know, from our earlier study using a TaqMan assay, that these gDNA samples have no detectable wildtype amplicons. Our explanation for this anomaly is that 3-GD of nearly pure mutant amplicons (Equation 12) generates Gaussians that overlap with those of mocks (Fig. 5A). In support of this hypothesis is our earlier finding that indels with sufficiently large insertions can mimic wildtype molecules in HRMA and constitute less than 10%. It is rather unlikely for mutant frequencies to approach such high levels in transient transfection experiments in the absence of drug selection. We therefore believe that this would not pose a significant hurdle for the GD model for estimation of editing efficiencies.

During GD of mocks, we were intrigued by the small discrepancy in the derivative melt curves at the melt transition temperature seen in single-Gaussian modeling. This seemed more pronounced in F8-S2 samples. We hypothesized that in F8-S2 amplicons, there were regions of the sequence that melted sooner or behaved as a nearly independent domain that was AT-rich. To identify these regions in the sequence, we wrote a Python function that determined the percentage of As and Ts in sliding windows of 10-mers that shifted by one nucleotide. The moving averages (period = 5) are shown in Fig. 8A and 8B (green traces). In the F8-S2 sequence, two initial broad regions with high AT content were visible (Fig. 8A). In contrast, in the CCR5 sequence, few AT-rich regions that seemed narrower were seen (Fig. 8 B).

**Fig. 8.**
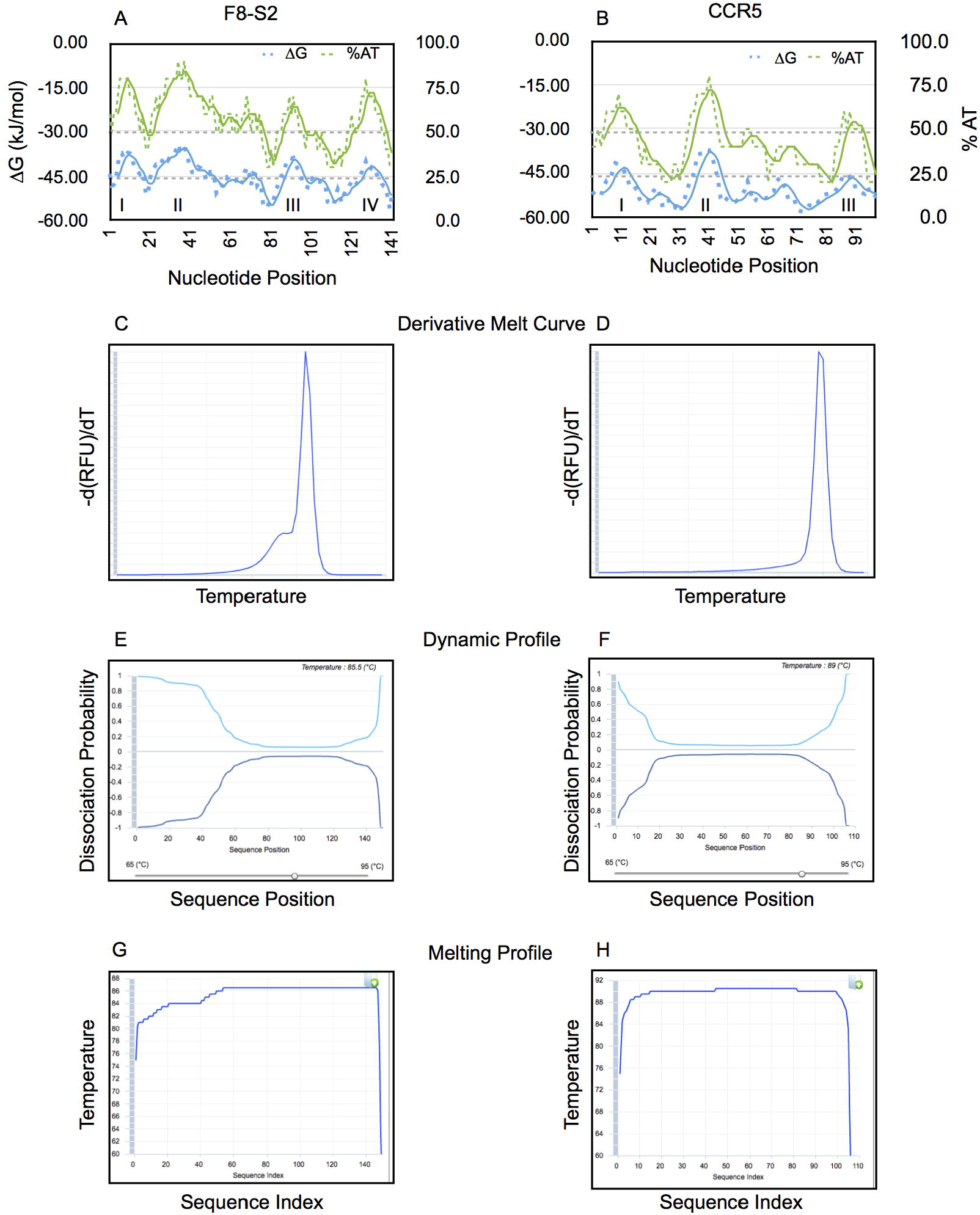
Analysis of F8-S2 and CCR5 target sequence features and melting properties *in silico*. Sliding window analysis of percentage of AT (%AT) in F8-S2 (A) or CCR5 (B) sequences of target sites amplified by PCR. The percentage of As and Ts were determined in a sliding overlapping window of 10-mers. The shift was by 1 bp. These are shown as green dashes. The data was smoothed using running averages with a period of 5 (solid green line). The sum of free energies (∆Gs) in a sliding window of 10-mers and a shift of 1 bp is shown along the left y-axis in kJ/mol (blue dots). The running averages were calculated as for %AT traces and are shown as blue traces. Putative AT-rich domains are marked I-IV. (C-H) The F8-S2 and CCR5 target sequences were used as input in the UMelt web analysis tool (29). UMelt predicted derivative melt curve (C and D), “Dynamic Profile” of melting (E and F) using a sliding temperature control that was situated close to the predicted Tm for each sequence to identify portions of the target sequences (nucleotide position indicated on the x-axis) that may have melted earlier than the rest. The web tool also provided a “Melting Profile” analysis that shows potential regions that might show greater tendency to melt earlier (G and H).

We wrote another Python function to compute the free energy of a 10-mer sequence window by using the nearest-neighbor method. For this analysis too, we used a sliding window that shifted by one nucleotide. The moving averages (period =2) are shown in Fig. 8 (blue traces). Again, the initial AT-rich region exhibited lower free energies (∆Gs) for F8-S2 sequence than that of the CCR5 sequence (Fig. 8A and 8B).

We next used the online web tool uMelt [29] to determine if the melting profiles of F8-S2 and CCR5 amplicon sequences could be distinguished by *in silico* analysis. For F8-S2 amplicon, the derivative melt curve predicted by uMelt web tool, showed a bulge in the early melt region (Fig. 8C). The Dynamic Profile window also predicted melting at earlier temperatures at both ends, particularly at the 5’ end of the sequence (Fig. 8D). The Melting Profile pane (Fig. 8E) also showed increased melting at lower temperatures for the first 50 base pairs. In contrast to F8-S2, for the CCR5 target sequence amplicon, the web tool predicted only a small deviation of melt curve in the early melt region (Fig. 8F). The Dynamic Profile (Fig. 8G) for CCR5 target amplicon also showed nearly equal rates of melting from both ends of the sequence with a barely visible enhancement for the left end. Likewise, the Melting Profile pane (Fig. 8H) showed very little propensity for a separate domain that exhibited different melting characteristics than the rest of the sequence for CCR5. The differences noted between the predicted derivative melt curves and the experimentally derived counterparts have been attributed to uMelt software being based on ∆Gs determined for pairs of nucleotides using a spectrophotometric method rather than on fluorescence emission from the binding of dsDNA-binding fluorophores. Nevertheless, uMelt analysis supports the two-Gaussian model for curve fitting of unmodified control samples.

In conclusion, this paper describes a method to correct high-resolution melt curves for temperature-dependent quenching of free and dsDNA-bound fluorophore. This is the first report, to the best of our knowledge, to demonstrate that first derivative melting curves of properly processed high-resolution melt curve data can be precisely modeled as a sum or superposition of Gaussian functions. The GD model successfully estimated efficiency of genome-editing by engineered sequence-directed endonucleases without a requirement for standard curves and has the additional advantage of being a single-tube method.

## Acknowledgments

We thank the Division of Hematology-Oncology, Bone Marrow Transplant and Cellular Therapy within the Department of Internal Medicine, Saint Louis University, for making available facilities and funding to carry out the described work. We also thank Mike Marcinkowski for editing the text and Matt Schuelke for reviewing the statistics used in evaluating GD models.

## Supporting information

Datasets containing Excel, Numbers (generated on Mac) and Jupyter Notebook files (for running Python programs used in the manuscript) can be accessed at: https://figshare.com/s/4f07b851af468f18b42d

